# Long-term impact of intensive postgraduate laboratory training at the Cold Spring Harbor Neurobiology of *Drosophila* summer course

**DOI:** 10.1101/369892

**Authors:** Sarah Ly, Karla Kaun, Chi-Hon Lee, David Stewart, Stefan R. Pulver, Alex C. Keene

## Abstract

Intensive postgraduate courses provide an opportunity for junior and senior level scientists to learn concepts and techniques that will advance their training and research programs. It is commonly assumed that short intensive courses have positive impacts within fields of research; however, these assumptions are rarely tested. Here we describe the framework of a long running postgraduate summer course at Cold Spring Harbor and attempt to quantify the impact made over its history. For over three decades, the *Drosophila* Neurobiology: Genes, Circuits & Behavior Summer Course at Cold Spring Harbor Laboratories (CSHL) has provided participants with intense instruction on a wide variety of topics and techniques in integrative neuroscience using *Drosophila* as a model organism. Students are introduced to the latest approaches for studying nervous system development, activity and connectivity, as well as complex behaviors and diseases. The course has a long history of successful alumni, many of whom describe participation in the course as foundational to their training. Student surveys of recent participants indicate a high level of satisfaction, improved career outcomes, and direct impact on publications. Analysis of student success reveals that over 64% of participants obtain independent faculty positions. Further, we describe ongoing efforts to enhance diversity and encourage access to scientific research at undergraduate-focused institutions. Together, our findings suggest that laboratory-intensive postgraduate courses provide a highly effective mechanism for scientific training that has lasting positive impacts on trainees.

## INTRODUCTION

Research on the fruit fly, *Drosophila melanogaster*, has played an important role in uncovering principles of nervous system structure and function. Fundamental insights include the first molecular descriptions of nervous system differentiation (reviewed in [1]), the identification and elucidation of axon guidance cues (reviewed in [2]), the first cloning of ion channels (reviewed in [3]), the first demonstration that SNARE proteins are required for chemical neurotransmission (reviewed in [4]), and the identification of single genes that regulate complex behavior (reviewed in [5–7]). More recent advances have been focused on neural circuit function and the neuronal basis for complex behavior (reviewed in [8]). *Drosophila* often leads the way in the development of genetic technologies for identifying, manipulating and monitoring neural circuits [9]. Furthermore, complex behaviors, including learning, decision-making, feeding, circadian rhythms, arousal/sleep, aggression, courtship, and addiction are now amenable to sophisticated experimental analysis. Increasingly, a comprehensive understanding of the functional connections between genes, proteins, neural circuits, and emergent behaviors is attainable [10–14]. Importantly, conservation of both the genes that underlie circuit formation and function as well as the information-processing logic of neural circuits means that discoveries and technologies developed in *Drosophila* are often readily translated into insights that directly inform research in mammalian systems [7, 15–17].

The growing diversity of genetic approaches and specialization of individual laboratories presents a notable impediment to scientific training. Creative and novel scientific approaches require a familiarity with cutting-edge genetic technology and approaches in model organisms, yet this type of specialized coursework is not provided in most laboratories or training programs. Laboratory-intensive courses provide trainees with rigorous experimental training that includes exposure to research areas and techniques that may not be available at their host institution. These field-specific courses are offered at institutions throughout the world, and are widely viewed as critical aspects of training for the scientific community. Although numerous descriptive reports have been written about the structure of various courses [18–21], understanding the true impact of these courses requires a quantitative assessment of course efficacy and student satisfaction that has yet to be conducted.

Max Delbruck organized the first modern course in Bacterial Viruses at Cold Spring Harbor Laboratory (CSHL). The objective was to create a large group of research workers of diverse backgrounds who were indoctrinated in the biological side of bacteriophage research and could work in a somewhat loose collaboration to explore this most promising field. The basic philosophy of this course included learning science by doing science, and of total immersion in the concepts and techniques of a new field. This philosophy has remained the cornerstone of CSHL postgraduate courses to the present date; its’ impact has been enormous, and has inspired fundamental concepts within the field of molecular biology. Modern neuroscience courses began at CSHL in 1981 with a course on Single Channel Recording, which was established shortly after Erwin Neher and Bert Sakmann first demonstrated the Nobel prize winning methodology.

In the 1970s, CSHL began offering several Neurobiology of *Drosophila* workshops that included contributions from many early leaders in the field including Bill Pak, Chun-Fang Wu, and Bob Horovitz. These workshops in the early days of *Drosophila* neurogenetics undoubtedly had an outsized role on the trajectory of the field and development of community that continues to endure today [22]. In 1984, the workshop was fully revamped to capture the incredible changes in the scientific landscape and was the birth of our present day — and longtime running — fly course. In 1984, an annual course focused on the nervous system of the fruit fly, *Drosophila melanogaster* was formalized. The course was offered under the direction of Ralph Greenspan, Lily Jan, Yuh-Nung Jan and Patrick O’Farrell. The course has now run without interruption for 35 years. Each year, approximately12 trainees are selected through a competitive admission process to participate in the three-week course, which is led by instructors from diverse backgrounds. Many of these trainees have gone on to become leaders in the field. Here, we describe the current course structure in the context of past course highlights and provide a systematic analysis of student satisfaction and the long-term career success of course participants. Our analysis reveals remarkable sustained achievement of course alumni, thus highlighting the potential of similar courses to catalyze scientific impact and promote diversity within the scientific community.

## METHODS

### Course Structure

The CSHL *Drosophila* neurobiology laboratory and lecture course is intended for researchers at all levels who want to use *Drosophila* as an experimental system to study the nervous system. Students of the course learn how to examine the larval and adult *Drosophila* nervous systems to study development, neurophysiology, neuroanatomy, and behavior. Students learn a wide range of methods and experimental approaches including genetic, electrophysiological, imaging and behavioral techniques. The course is actively designed to balance in-depth training with an expanded breadth of topics. Daily seminars introduce the history behind special topics in *Drosophila* research while providing updated current knowledge by expounding upon recent contributions to the literature and generating interactive discussions about outstanding questions in the field. Student discussion is actively encouraged and major emphasis is placed on experimental methodology. Guest lecturers provide original preparations for demonstration, discussion, and direct laboratory experimentation in their areas of special interest and expertise. Each lecturer provides a supplementary reading list and several background papers or reviews that students are encouraged to read before each lecture. Students are also provided with detailed protocols for all laboratory experiments to facilitate the direct transfer of their cutting-edge training in the course to their home labs. If requested, students can also take novel genetic and molecular reagents that are rapidly advancing this field back with them to their own laboratories.

The material provided by instructors in lectures and laboratory exercises is supplemented by evening seminars that provide specific information on the current status of research in the invited speakers’ area of expertise. Speakers often spend many hours informally discussing the participants’ experiments at the course in relation to their current research interests. Daily formal and informal discussions between students, faculty, and invited speakers on the finer points of the techniques and concepts being taught in each course are a valuable source of intellectual stimulation. These discussions also allow the instructors and lecturers to provide insight and advice on how to address the specific problems being encountered by students in their own research at home. These facets of the program all contribute to the total immersion of the participants in the subject. The presence of visiting experts for each area of the course allow for technical training at a depth that is not possible in other courses and workshops. Participants are able to focus their attention on the scope of the course without the distraction of other responsibilities, promoting full immersion in the relevant methodologies and concepts.

The daily schedule of the *Drosophila* Neurobiology Course is designed such that students spend most of their time performing hands-on laboratory activities to consolidate information they learned during seminars about cutting-edge research and lectures of basic neurobiological, genetic, developmental, molecular and behavioral concepts. Typically, students receive two lectures in the morning: one on basic concepts, followed by one on research techniques. Students are subsequently in the lab all afternoon and into the evening. Approximately half of the evenings of the course have additional lectures designed to expose students to exciting contemporary research in *Drosophila* neurobiology, and provide perspective on how current work complements research in other model organisms. The overall three-week schedule is formulated so that students first learn the basic neurodevelopmental and genetic concepts prior to learning physiology, sensory systems, simple behavior, and finally complex behavior. The twelve students accepted to the course are encouraged to think independently and use cooperative learning to maximize their own and one another’s learning.

A central goal of the course is to expose students to techniques that can be readily implemented at their home institution. The course covers a broad range of techniques, such as Ca^2+^ imaging, neuromuscular junction electrophysiology, and behavioral analysis using cutting-edge genetic tools. More recently, Do-it-yourself (DIY) methodologies, such as 3D printing for constructing behavioral systems, has encouraged creative approaches for understanding the circuits and molecules that regulate physiology and behavior [23, 24]. The potential for students to establish similar DIY systems at their home institutions is emphasized, thereby increasing the impact of these techniques taught at the course.

### Admissions and recruitment

Applications are open to candidates at any stage in their postgraduate career from academia and industry within the United States and from overseas. Participants are selected by the course instructors on the basis of several criteria including quality of proposed research, demonstrated need for training in a specific research area, institutional/community impact of training, national versus international reach, US underrepresented minority (URM) status, breadth of representation of scientific approaches, and gender balance. Faculty make efforts to select multiple stages of trainees, ranging from graduate students to senior investigators. This distribution offers several distinct advantages: students and fellows often benefit from working closely with more mature advanced scientists in an informal setting, particularly in terms of learning approaches and priorities. These interactions have consistently generated a dynamic and stimulating environment that facilitates peer-to-peer mentorship and cohort cohesion that is central to the objectives of the course.

### Curriculum selection and evaluation

The course curriculum is determined by the course instructors, and progress is monitored by the senior administrative staff through discussion and evaluation during and after each course with faculty, students, and guest lecturers (including former instructors). Over the decades of the course, the curriculum and format have changed dramatically, often to reflect the current focus of the field and state-or-the-art genetic technology. For example, the initial years of the course were intensely focused on genetic mapping, consistent with the common approach of mutagenesis-based screens in the 1980’s [25]. This then changed during the early 2000’s into a format with separate teaching modules on development, physiology, and behavior. More recently, the format has shifted to integrative laboratories that are not separated by research area. There has also been a recent emphasis on circuit dissection and implementation of state-of-the-art genetic tools for manipulating gene expression and neuronal function.

### Data collection and outcomes analysis

Each year, students complete anonymous course evaluation forms at the end of the course with the primary aim of using the feedback to continue improving the quality of the course. The course undoubtedly impacts other participants, including instructors and teaching assistants. While impact and satisfaction assessment of these groups would be highly valuable, the metrics for determining these aspects is complex, and much of these data have not been collected. Therefore, our analysis focuses on outcomes for trainees. All data collection, analysis and presentation was performed in accordance with Cold Spring Harbor Laboratories IRB Protocol 17-019.

## RESULTS AND DISCUSSION

### Informal assessment of the course

To a large degree, queries of course impact come from informal interactions or feedback requests to former participants. These queries are undoubtedly biased, as the requests are largely made to senior faculty who have remained in the field. However, the degree to which many of these former students credit the class with playing a critical role in their education was notable. For example, Claude Desplan (Class of ’85) stated, ‘This was a very long time ago, but I sincerely think that it has changed my perception of the world of Science.’ In addition, nearly all respondents described how the course provided the unique opportunity to interact with leading senior scientists. Leslie Voshall (Class of ’91) stated, ‘The opportunity to mingle with world-class scientists in an informal setting was matchless. That year, I had the chance to interact with Seymour Benzer, who provided major input into my thesis work on biological clocks.’ Finally, respondents described how the course developed a sense of community that included students and encouraged scientific creativity. Joel Levine (Class of ’95) stated ‘it left me with a sense that I was part of a culture and that almost anything is possible with the fly.’

### Course Participants

Admission to the course is competitive, with an average acceptance rate of approximately 30% over the last nine years. From 2008-2016, the course typically had 12 students from diverse career stages (Table 1). The majority of students (63%) were doctoral trainees, with an additional 27% postdoctoral trainees and 9.4% senior scientists. The course has also historically maintained gender and geographic diversity, with a nearly equal male:female ratio, and 32% of students coming from laboratories that are outside of the United States.

**Table 1:** Makeup of course students, assistants and faculty from 2008-2016. The number instructors (Course organizers), lecturers, teaching assistants, applications, and students for each year from 2008-2016. The breakdown of students (bold) between grad student, postdoc, and PI/senior scientist, as well as gender. The percentage of students from the US. URM denotes US underrepresented minority.

|  | <u>2008</u> | <u>2009</u> | <u>2010</u> | <u>2011</u> | <u>2012</u> | <u>2013</u> | <u>2014</u> | <u>2015</u> | <u>2016</u> | **<u>TOTAL</u>** |
| --- | --- | --- | --- | --- | --- | --- | --- | --- | --- | --- |
| Instructors | 4 | 3 | 3 | 3 | 3 | 3 | 3 | 3 | 3 |  |
| Lecturers | 29 | 21 | 17 | 21 | 14 | 18 | 19 | 22 | 26 |  |
| Assistants | 5 | 5 | 6 | 7 | 4 | 4 | 3 | 3 | 10 |  |
| Applicants | 52 | 34 | 24 | 23 | 43 | 42 | 32 | 32 | 36 | **319** |
| **Students** | **11** | **12** | **12** | **11** | **12** | **12** | **12** | **12** | **12** | **106** |
| Grad Stud | 7 | 6 | 7 | 8 | 8 | 10 | 9 | 6 | 6 | **67** |
| Postdoc | 2 | 3 | 4 | 3 | 3 | 2 | 3 | 4 | 5 | **29** |
| PI/Sen Sci | 2 | 3 | 1 | - | 1 | - | - | 2 | 1 | **10** |
| Gender | 6m/5f | 5m/7f | 6m/6f | 5m/6f | 6m/6f | 5m/7f | 7m/5f | 6m/6f | 5m/7f | **51m/55f** |
| US (%) | 7(64%) | 9(75%) | 6(50%) | 6(55%) | 9(75%) | 9(75%) | 8(67%) | 9(75%) | 9(75%) | **72(68%)** |
| URM | 1 | 1 | 1 | 1 | 1 | 2 | na | 2 | 3 | **12(17%)** |

### Instructors and Lectures

The course faculty is comprised of researchers who are active in fly genetics, neurodevelopment, physiology, and behavior. The course is led by three to four instructors (organizers), who collaboratively design the course syllabus, plan the course schedule, and invite suitable lecturers based on current *Drosophila* neurobiology research and student feedback. Each year between 2008and 2011, approximately sixteen lecturers gave both seminars and laboratory sessions, whereas an additional ~eight lecturers gave evening seminars on cutting-edge research in their laboratories. Lecturers formed a cross section of the *Drosophila* community, coming from the institutions in the United States (83.6%) and outside the US (16.4%). They represent primarily research universities (61.2%) and research institutes (37.7%), but also government (0.9%) and teaching colleges (0.9%). A range of career stages are represented from full professors (12.93%) and tenure track faculty (58.7%) to postdoctoral fellows graduate students and technicians (33.6%). Gender balance was actively encouraged in recruitment of course lecturers, instructors and TAs, with an overall 60.3% male:39.7% female ratio across the 8 years analyzed.

### Student Satisfaction

The perceived impact of the course on students was quantified through a standardized questionnaire solicited each year from 2012-2016. The survey was designed to measure both personal satisfaction and the perceived contribution of the course to the trainees’ scientific career. Evaluation sheets were typically circulated on the last day of the course and students were encouraged (but not required) to complete and return the evaluations before they departed from CSHL. Importantly, evaluations were handwritten and completely anonymous. Each evaluation included a scoring system in response to questions about different aspects of the course: 1 (needs improvement) - 5 (exceptional). Table S1 includes a summary of average scores for the course since 2012. Evaluations reveal a clear level of satisfaction amongst each student class upon completion of the course. While enthusiasm for the course and points of strength vary from year to year, students expressed strong overall enthusiasm for all aspects of the course, with all areas averaging a score of 4.1 (out of 5) or greater (Table 2). On average, the highest scores were obtained for helpfulness of instructors, receiving an average score of 4.88.

**Table 2:** Student course evaluations from 2012-2016. Average scores from course evaluations over a five-year period. Scale: 1-Poor->5-Excellent.

|  | <u>**2012**</u> | <u>**2013**</u> | <u>**2014**</u> | <u>**2015**</u> | <u>**2016**</u> |
| --- | --- | --- | --- | --- | --- |
| In general, did the course meet your needs /expectations? | 4.5 | 4.6 | 4.1 | 4.8 | 4.9 |
| Were the lecture topics well chosen? | 4.5 | 4.5 | 4.1 | 5.0 | 4.8 |
| Was the level of the lectures appropriate? | 4.3 | 4.7 | 4.5 | 4.8 | 4.8 |
| Were the presentations clear? | 4.2 | 4.5 | 4.1 | 4.5 | 4.6 |
| Were the instructors helpful? | 4.9 | 4.9 | 4.6 | 5.0 | 5.0 |
| Was the selection of lab exercises appropriate? | 4.1 | 4.5 | 3.6 | 4.7 | 4.8 |
| Was there sufficient/too much supervision of the lab? | 4.3 | 4.5 | 4.2 | 4.5 | 4.8 |
| Were the labs well enough equipped? | 3.9 | 4.4 | 4.2 | 4.3 | 4.8 |
| What was the utility and quality of the written protocols? | 3.8 | 4.4 | 3.4 | 4.3 | 4.5 |
| How was the course work load? | 4.5 | 4.3 | 4.4 | 4.4 | 4.8 |

### Long-term Qualitative Impacts

In total, more than 300 trainees have participated in the course over its 36 year history. We assessed longer-term impacts by 1) soliciting feedback from alumni in previous years of the course and 2) tracking of careers of former students.

We solicited feedback from alumni from 1996 to 2015 and asked how the course contributed to the individual’s intellectual development, technical expertise, publications, as well as impact on the field as a whole. In all, 85 alumni responded in full to the survey. The results of this survey reveal that the majority of students found the course to be helpful for their career (87%; Table 2). In addition, the vast majority reported that the course helped initiate new research directions (78%; Table 3), and was a highlight of their scientific career (95%). The high satisfaction with the course is indicative of substantive and long-lasting positive impact on career development.

**Table 3.** Long-term impact assessment of students from 1996-2015. Previous students were contacted and asked about the impact the course had on the career over the summer of 2016. The number of responses for the 85 recipients for each category are listed.

|  | Strongly Agree | Agree | Neutral | Disagree | Strongly Disagree |
| --- | --- | --- | --- | --- | --- |
| The course helped me achieve my next career transition. | 41 | 31 | 11 | 0 | 1 |
| The course had a positive impact on my scientific career as a whole. | 71 | 13 | 1 | 0 | 0 |
| The course helped me initiate a new research direction. | 39 | 29 | 15 | 1 | 1 |
| The course taught me new techniques that were subsequently applicable to my research | 56 | 24 | 2 | 0 | 2 |
| The course was a highlight in my scientific career. | 56 | 24 | 2 | 0 | 2 |
| The course has a positive impact on the broader biological research community. | 62 | 19 | 3 | 1 | 0 |
|  | YES | NO |  |  |  |
| Do you still work in this scientific area? | 70 | 12 |  |  |  |
|  | 10+ | ~7 | ~4 | ~2 | 0 |
| Number of research publications that could be attributed to taking this course? | 4 | 10 | 14 | 49 | 7 |
|  | 3+ | 3 | 2 | 1 | 0 |
| Number of patents that could be attributed to taking this course? | 1 | 0 | 1 | 3 | 72 |

The success of the course can also be measured by assessing the career trajectories of alumni. An exhaustive internet search of publicly available information was conducted to identify the current positions of course graduates from 1983-2016 (Fig 1). We were able to confirm the career status of 287 former students of the course. This revealed that 64% are currently in academic faculty positions with 55% of identified course alumni in tenure track positions. This number is remarkably high compared to recent estimates of success in obtaining an academic faculty position. In the United States, an estimated 65% of graduate students in biomedical fields move on to post-docs, and only 10-13% secure faculty positions [26]. In this course, a relatively small percentage of graduates (29 students, 9%) obtained industry research positions, likely indicative of the course focus on *Drosophila* basic science research and student success in obtaining tenure track positions. Importantly, alumni also showed a diverse set of non-research careers; examples included scientific journal editors, teachers, and entrepreneurs. The long-term impact on student success is indicative of a high return on investment, both in terms of student time and scientific funding. Further, while difficult to quantify, it is likely that the course’s effectiveness extends beyond alumni as they were likely to share their course training with individuals in their local scientific communities. The fact that a large number of respondents indicated that publications could be attributed to taking the course suggests that there was indeed transfer of knowledge to local communities that was effectively translated into observable research outputs. To this point, 90% of respondents confirmed that at least 2 publications could be attributed to taking the course (Table 4).

**Figure 1.**
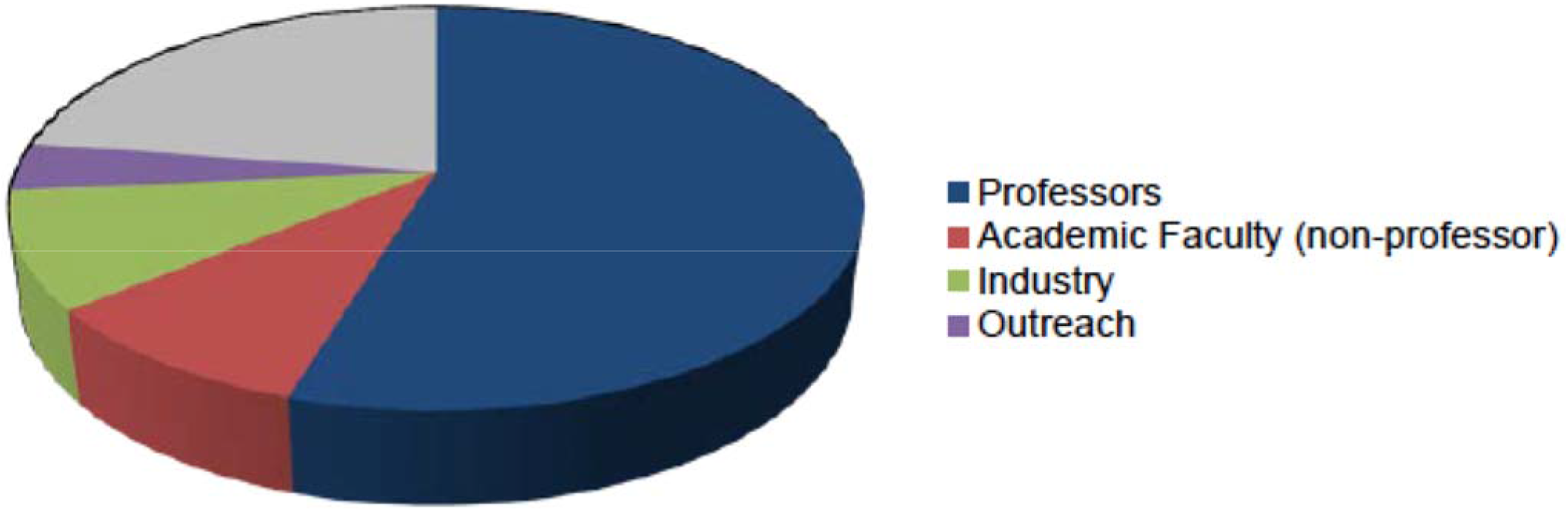
Current professional status of course graduates. In total, 55% of eligible course alumni received professor appointments, while 64% hold academic appointments (including non-professor appointments). 9% of alumni work in industry, 4% in outreach. Grey (23%) represents other professional positions.

Beyond assisting the careers of individual students, the course is uniquely positioned to provide training opportunities to those with limited access to research resources. While many training opportunities - including National Institutes of Health NRSA fellowships - strongly consider training environment, this course focuses on providing access to students from diverse geographic and scientific backgrounds. The ability to recruit and train students from a wide range of scientific and cultural backgrounds has the potential to extend neuroscience training to communities that are underrepresented in scientific funding. From 2008 until the present, the course has steadily increased the numbers of students from URM backgrounds (Table 1). In recent years (2015-present), special efforts have been made to recruit students and lecturers who teach at URM serving institutions. This approach provides the course with a way to extend Drosophila neuroscience into underserved communities and maximize the social and scientific impact of the course.

### The role of selection bias in outcomes analysis

Although the data obtained from previous course participants clearly indicates that the course is associated with future success in academia, we acknowledge that biases likely contribute to this association. For example, the applicants to this program may be from a specific pool of researchers who have knowledge of the course and are motivated to apply to a course, which removes them from their research for nearly an entire month in the summer. Moreover, although careful consideration is given to ensuring the participants chosen to attend the course are balanced in terms of geographical, scientific and cultural diversity, selection of the candidates may favor researchers who would have been successful in the absence of taking the course. At this time, it is not possible to determine how these biases contribute to the success of the course participants. However, future analysis that compares course participants to those not selected to participate in the course may elucidate some of these concerns. Nevertheless, the analysis highlighting long-term student success clearly makes it likely that connections established during the course have a lasting impact on the scientific community.

### Outlook for the future

The CSHL *Drosophila* Neurobiology course has demonstrated remarkable success over more than three decades. This likely reflects both a general effectiveness of the intensive short summer course format for postgraduate researchers as well as the specific approaches utilized by this course. In particular, the flexible curriculum defined by rotating organizers, instructors, and laboratories, has ensured that cutting-edge research and techniques are consistently incorporated over the multi-decade duration of the course. Moving forward, we plan to continue this innovation by investing in growing areas of research. For example, future iterations of the course are likely to emphasize computational approaches to neuroscience, functional imaging, automated behavioral data acquisition, and quantitative analysis of animal behavior. In addition, we will increase the emphasis on DIY approaches to encourage students to develop creative solutions to investigate the mechanisms that underlie brain physiology and behavior.

Finally, we hope to leverage the success of this course to extend beyond the reach of CSHL. For example, forming collaborations with faculty outside CSHL to extend course laboratory modules into undergraduate curriculums will significantly broaden and enhance the impact of the course. Indeed, recent participants at the course have adapted CSHL modules for use in undergraduate education and these modules are now published and freely available [27–29]. Finally, generation of publicly accessible protocoxl videos will enhance the broader impact of this course. These points of emphasis, combined with incorporating rapidly improving genetic technology in the fly, will allow this course to remain current and innovative into the future.

### Conclusions

*Drosophila melanogaster* is uniquely suited to teach fundamental principles in neuroscience research. The amenability of the model to laboratory manipulation and its short generation time allow for the efficient implementation of cutting-edge genetic technology for investigating development, physiology, and behavior in a short-course format. This manuscript provides an overview of the course curriculum and structure. In addition, we provide quantitative analyses which reveal that the *Drosophila* Neurobiology Course has experienced remarkable success in terms of student satisfaction and career outcomes. The active emphasis on selecting students from diverse backgrounds further promotes greater scientific access in the field, gender equality, and success for underrepresented minorities. Thus, our findings suggest that courses such as the one described here have the ability to dramatically encourage scientific career success, broader dissemination of cutting-edge research, and positive social impact in the global science community.

## Acknowledgments

We would like to thank the CSHL staff, and the students, lecturers and teaching assistants that participated in the *Drosophila* Neurobiology Course. We would also like to extend a special thank-you to past course instructors Kate O’Connor-Giles (Brown University), Adrian Rothenfluh (University of Utah) and Greg Macleod (Florida Atlantic University) for assistance with an early version of portions of this text. This work was funded by the National Science Foundation Division of Integrative Organismal Systems (NSF IOS 1523125) and National Institute for Drug Abuse (NIDA R13DA034437) and NSF awards 165674 and 146465 to ACK.

## Works Cited

1. Kohwi M, Doe CQ (2013) Temporal fate specification and neural progenitor competence during development. Nat Rev Neurosci 14:823–38. doi: 10.1038/nrn3618

2. Evans TA (2016) Embryonic axon guidance: insights from Drosophila and other insects. Curr Opin Insect Sci 18:11–16. doi: 10.1016/j.cois.2016.08.007

3. Frolov R V, Bagati A, Casino B, Singh S (2012) Potassium channels in Drosophila: historical breakthroughs, significance, and perspectives. J Neurogenet 26:275–290. doi: 10.3109/01677063.2012.744990

4. Harris KP, Littleton JT (2015) Transmission, development, and plasticity of synapses. Genetics 201:345–375. doi: 10.1534/genetics.115.176529

5. Sokolowski MB (2001) Drosophila: genetics meets behaviour. Nat Rev Genet 2:879–890.

6. Vosshall LB (2007) Into the mind of a fly. Nature 450:193–197. doi: 10.1038/nature06335

7. Bellen HJ, Tong C, Tsuda H (2010) 100 years of Drosophila research and its impact on vertebrate neuroscience: a history lesson for the future. Nat Rev Neurosci 11:514–22. doi: 10.1038/nrn2839

8. Kazama H, Wilson RI (2008) Homeostatic matching and nonlinear amplification at identified central synapses. Neuron 58:401–413. doi: S0896-6273(08)00184-0[pii]10.1016/j.neuron.2008.02.030

9. Venken KJ, Simpson JH, Bellen HJ (2011) Genetic manipulation of genes and cells in the nervous system of the fruit fly. Neuron 72:202–230. doi: 10.1016/j.neuron.2011.09.021

10. Zwart MF, Pulver SR, Truman JW, et al. (2015) Selective Inhibition Mediates the Sequential Recruitment of Motor Pools. Neuron 91:615–628. doi: 10.1016/j.neuron.2016.06.031

11. Aso Y, Sitaraman D, Ichinose T, et al. (2014) Mushroom body output neurons encode valence and guide memory-based action selection in Drosophila. Elife 3:e04580. doi: 10.7554/eLife.04580

12. Seidner G, Robinson JE, Wu M, et al. (2015) Identification of Neurons with a Privileged Role in Sleep Homeostasis in Drosophila melanogaster. Curr Biol 25:2928–2938. doi: 10.1016/j.cub.2015.10.006

13. Vogelstein JT, Park Y, Ohyama T, et al. (2014) Discovery of brainwide neural-behavioral maps via multiscale unsupervised structure learning. Science (80-) 344:386–392. doi: 10.1126/science.1250298

14. Klein M, Afonso B, Vonner AJ, et al. (2015) Sensory determinants of behavioral dynamics in Drosophila thermotaxis. Proc Natl Acad Sci U S A 112:E220–9. doi: 10.1073/pnas.1416212112

15. Stavropoulos N, Young MW (2011) Insomniac and cullin-3 regulate sleep and wakefulness in drosophila. Neuron 72:964–976. doi: 10.1016/j.neuron.2011.12.003

16. Seelig JD, Jayaraman V (2015) Neural dynamics for landmark orientation and angular path integration. Nature 521:186–191. doi: 10.1038/nature14446

17. Pulver SR, Griffith LC (2010) Spike integration and cellular memory in a rhythmic network from Na(+)/K(+) pump current dynamics. Nat Neurosci 13:53–59. doi: nn.2444[pii]10.1038/nn.2444

18. Patel VL, Branch T, Cimino A, et al. (2005) Participant perceptions of the influences of the NLM-sponsored woods hole medical informatics course. J Am Med Informatics Assoc. doi: 10.1197/jamia.M1662

19. Davidson C, Creson D (1971) A summer program for minority students in a medical setting--a progress report. Tex Rep Biol Med 29:443–50.

20. Fallon JF (2002) How serendipity shaped a life: An interview with John W. Saunders, Jr. Int. J. Dev. Biol.

21. Bridges J, Miller CJ, Kipnis DG (2006) Librarians in the Woods Hole Biomedical Informatics Course. Med Ref Serv Q. doi: 10.1300/j115v25n01_07

22. Jan LY, Jan YN (2018) Influences: Cold Spring Harbor summer courses and Drosophila melanogaster neurogenetics. J Gen Physiol 150:733–735. doi: 10.1085/jgp.201812063

23. Baden T, Chagas AM, Gage G, et al. (2015) Open Labware: 3-D Printing Your Own Lab Equipment. PLoS Biol. doi: 10.1371/journal.pbio.1002086

24. Geissmann Q, Rodriguez LG, Beckwith EJ, et al. (2017) Ethoscopes: An Open Platform For High-Throughput Ethomics. bioRxiv 113647. doi: 10.1101/113647

25. St Johnston D (2002) The art and design of genetic screens: Drosophila melanogaster. Nat Rev Genet 3:766–778. doi: 10.1038/nrg751

26. Powell K (2015) The future of the postdoc. Nature 520:144–147. doi: 10.1136/vr.117.21.563

27. McKellar Claire, E; Wyttenbach, Robert A (2017) A Protocol Demonstrating 60 Different Drosophila Behaviors in One Assay. J Undergrad Neurosci Educ 15:A110–A116.

28. Titlow JS, Johnson BR, Pulver SR (2015) Light Activated Escape Circuits: A Behavior and Neurophysiology Lab Module using Drosophila Optogenetics. J Undergr Neurosci Educ 13:A166–73.

29. Scaplen KM, Mei NJ, Bounds HA, et al. (2019) Automated real-time quantification of group locomotor activity in Drosophila melanogaster. Sci Rep. doi: 10.1038/s41598-019-40952-5

